# Accelerating COVID-19 research with graph mining and transformer-based learning

**DOI:** 10.1101/2021.02.11.430789

**Authors:** Ilya Tyagin, Ankit Kulshrestha, Justin Sybrandt, Krish Matta, Michael Shtutman, Ilya Safro

**Affiliations:** Center for Bioinformatics and Computational Biology, University of Delaware, Newark, DE; Computer and Information Sciences, University of Delaware, Newark, DE; School of Computing, Clemson University, Clemson, SC; Charter School of Wilmington, Wilmington, DE; Drug Discovery and Biomedical Sciences, University of S. Carolina, Columbia, SC

**Keywords:** Hypothesis Generation, Literature-Based Discovery, Transformer Models, Semantic Networks, Biomedical Recommendation

## Abstract

In 2020, the White House released the, “Call to Action to the Tech Community on New Machine Readable COVID-19 Dataset,” wherein artificial intelligence experts are asked to collect data and develop text mining techniques that can help the science community answer high-priority scientific questions related to COVID-19. The Allen Institute for AI and collaborators announced the availability of a rapidly growing open dataset of publications, the COVID-19 Open Research Dataset (CORD-19). As the pace of research accelerates, biomedical scientists struggle to stay current. To expedite their investigations, scientists leverage hypothesis generation systems, which can automatically inspect published papers to discover novel implicit connections. We present an automated general purpose hypothesis generation systems AGATHA-C and AGATHA-GP for COVID-19 research. The systems are based on graph-mining and the transformer model. The systems are massively validated using retrospective information rediscovery and proactive analysis involving human-in-the-loop expert analysis. Both systems achieve high-quality predictions across domains (in some domains up to 0.97% ROC AUC) in fast computational time and are released to the broad scientific community to accelerate biomedical research. In addition, by performing the domain expert curated study, we show that the systems are able to discover on-going research findings such as the relationship between COVID-19 and oxytocin hormone.

**Reproducibility:** All code, details, and pre-trained models are available at https://github.com/IlyaTyagin/AGATHA-C-GP

**CCS CONCEPTS:** • **Applied computing** → *Bioinformatics*; *Document management and text processing*; • **Computing methodologies** → *Learning latent representations*; *Neural networks*; **Information extraction**; **Semantic networks**.

## 1 INTRODUCTION

Development of vaccines for COVID-19 is a major triumph of modern medicine and humankind’s ability to accelerate scientific research. While we are all hoping to see large-scale positive changes from fast mass adoption of the existing vaccines, there remain significant open research questions around COVID-19. The scientific community has a responsibility to do everything possible to block the ongoing transmission of the dangerous virus and *accelerate research to mitigate its consequences*. We present the following automated knowledge discovery system in order to propose new tools that could compliment the existing arsenal of techniques to accelerate biomedical and drug discovery research for events like COVID-19.

The COVID-19 pandemic became one of the most important events in the information space since the end of 2019. The pace of published scientific information is unprecedented and spans all resolutions, from the news and pop-science articles to drug design at the molecular level. The pace of scientific research has already been a significant problem in science for years [29], and under current circumstances this factor becomes even more pronounced. Several thousands papers are being added *weekly* to CORD-19 [39] (the dataset of publications related to COVID-19) and even more in MEDLINE [1]. As a result, groups working on similar problems may not be immediately aware of the other’s findings, which can lead to inefficient investments and production delays.

Under normal circumstances, the MEDLINE database of biomedical citations receives approximately 950,000 new papers per year. Currently this database indexes 31 million total citations. This pace challenges traditional research methods, which often rely on human intuition when searching for relevant information. As a result, the demand for modern AI solutions to help with the automated analysis of scientific information is incredibly high. For instance, the field of drug discovery has explored a range of AI analytical tools to expedite new treatments [12]. Designing lab experiments and finding candidate chemical compounds is a costly and long-lasting procedure, often taking years. To accelerate scientific discovery, researchers came up with a family of strategies to utilize public knowledge from databases like MEDLINE that are available through the National Institute of Health (NIH), which facilitate *automated hypothesis generation* (HG) also known as literature-based discovery. Undiscovered public knowledge, information that is *implicitly* present within available literature, but is not yet *explicitly* known by an individual who can act on that information, represents the target of our work.

Although, there are quite a few automated HG systems [12] including those we have previously proposed [35, 37], *none of them is currently customized and available in the open domain to massively process COVID-19 related queries*. In addition to the traditional general requirements for HG systems, such as high-quality results of hypotheses, interpretability and availability for broad scientific community, a specific demand for COVID-19 data analysis requires: (1) customization of the vocabulary and other logical units such as subject-verb-object predicates; (2) customization of the training data that in the reality of urgent research contains a lot of controversial and incorrect information; (3) models for different information resolutions; and (4) validation on the on-going domain-specific discovery.

### Our contribution

In this work we bridge this gap by releasing, AGATHA-C and AGATHA-GP, reliable and easy to use HG systems that demonstrate state-of-the art performance and validate their inference capabilities on both COVID-19 related and general biomedical data. To make them closely related to different goals of COVID-19 research, they correspond to micro(AGATHA-C, for COVID-19) and macroscopic (AGATHA-GP, for general purpose) scales of knowledge discovery. Both systems are able to process any queries to connect biomedical concepts but AGATHA-C exhibits better results on the molecular scale queries, e.g., those that are relevant to drug design, and AGATHA-GP works better for general queries, e.g., establishing connections between certain profession and COVID-19 transmission.

Both systems are the next generation of the AGATHA knowledge network mining transformer model [37]. They substantially improve the quality of the previous AGATHA by introducing new information layer into multi-layered semantic knowledge network pipeline, and expanding new information retrieval techniques that facilitate inference. We deploy the deep learning transfer model trained with up-to date datasets and provide easy to use interface to broad scientific community to conduct COVID-19 research. We validate the system via candidate ranking [36, 37] using very recent scientific publications containing findings absent in the training set. While the original AGATHA has demonstrated state-of-the-art performance for the time of its release, AGATHA and other systems were found to perform with notably lower quality on extremely rapidly changing COVID-19 research. We demonstrate a remarkable improvement in the range of approximately 20-30% (in ROC-AUC) on the average on different types of queries with very fast query process that allows massive validation. In addition, we demonstrate that the proposed system can identify recently uncovered gene (BST2) and hormone (oxytocin and melatonin) relationships to COVID-19, using only papers published before these connections were discovered.

### Reproducibility

All code, details, and pre-trained models are available at https://github.com/IlyaTyagin/AGATHA-C-GP

## 2 BACKGROUND

**CORD-19 dataset** [39] was released as a response to the world’s COVID-19 pandemic to help data science experts and researchers to tackle the challenge of answering the high priority scientific questions. It updates daily and was created by the Allen Institute for AI in collaboration with Microsoft Research, NLM, IBM and other organizations. At the time of this publication it contains over 400.000 scientific abstracts and over 150.000 full-text papers about coronaviruses, primarily COVID-19.

**MEDLINE** is a database of NIH that includes almost 31 million citations (as of 2021) of scientific papers related to the biomedical and related fields. Some of the citations are provided with MeSH (Medical Subject Headings) terms and other metadata. MEDLINE is one of the largest and well-known resources for biomedical text mining.

### Hypothesis Generation Systems

The HG field has been present in information sciences for several decades. The first notable approach was proposed by Swanson et al. in 1986 [33], which is called the A-B-C model. The concept of A-B-C model is to discover intermediate (B) terms which occur in titles of publications for both terms A (source) and C (target). In their experiments, Swanson et al. discovered an implicit connection between Raynauld’s syndrome (term A) and fish oil (term C) through blood viscosity (term B), which was mentioned in both sets. The hypothesis that fish oil can be used for patients with Raynaud’s disease was experimentally confirmed several years later [10]. The key idea of the proposed method is that all fragmented bits of information are explicitly known, but their implicit relationships is what HG systems are aimed to uncover.

We note the difference between HG and traditional information retrieval. The information retrieval techniques which represent the vast majority of biomedical literature based discovery systems are trained and (what is even more important) validated to retrieve *existing information* whereas the HG techniques predict *undiscovered knowledge* and thus must be massively validated on it. The HG validation requires training the system strictly on historical data rather than sampling it over the entire time.

The advances in machine and deep learning transformed the algorithmics of HG systems (see Sec. 9) that are now able to process much larger information volumes demonstrating much higher quality predictions. However, lack of broader applicability of HG systems in the situation with COVID-19 pandemic demonstrates that several major issues exist and require immediate attention:

1. Most of the existing HG systems are domain-specific (e.g., gene-disease interactions) that is usually expressed in limiting the processed information (e.g., significant filtering vocabulary and papers to a specific domain in probabilistic topic modeling [38]);
2. A proper validation of HG system remains a technical problem because multiple large-scale models have to trained with all heterogeneous data carefully eliminated several years back;
3. Moreover, a large number of HG systems are not massively validated at all except of very old findings rediscovery [28] or demonstrating of just a few proactive examples in humanly curated investigation; and
4. Interpretability and explainbability of generated hypotheses remains a major issue.

**The UMLS Metathesaurus** [7] is the NIH database containing information about millions of concepts (both medical and general) and their synonyms. Metathesaurus accumulates information about its entries from more than 200 different vocabularies allowing to map and connect concepts from different terminologies. Metathesaurus also keeps metadata about the concepts such as semantic types and their hierarchy. The core unit of information in UMLS is the concept unique identifier, or CUI. CUI is a codified representation of a specific term, which includes its different atoms (spelling variants or translations of the term on other languages), vocabulary entries, definitions and other metadata.

**SemRep** [4] is a software kit developed by NIH for extraction of semantic predicates (subject-verb-object triples) from the provided corpus. It also allows to extract entities not involved in any semantic predicate, if the corresponding option is selected. The official example of possible SemRep output is: INPUT = “We used hemofiltration to treat a patient with digoxin overdose that was complicated by refractory hyperkalemia.”, OUTPUT = “Hemofiltration-TREATS-Patients; Digoxin overdose-PROCESS_OF-Patients; hyperkalemia-COMPLICATES-Digoxin overdose; Hemofiltration-TREATS(INFER)-Digoxin overdose”. SemRep handles word sense disambiguation and performs terms mapping to the corresponding CUIs from UMLS metathesaurus.

**ScispaCy** [24] ScispaCy is a special version of spaCy maintained by AllenAI, containing spaCy models for processing scientific and bio-related texts. ScispaCy models are trained on different sources, such as PMC-pretrained word2vec representations, MedMentions Entity linking Dataset and so on. SciSpacy can handle various NLP tasks, such as NER, dependency parsing and POS-tagging, where achieves state of the art performance.

**SciBERT** [6] is a BERT-like transformer pretrained language model, where full-text scientific papers were used as a training dataset. Embeddings are learned in a word-piece fashion, which makes them capture the relationships between not only words in a sentence, but also between word parts in each word.

**FAISS** [15] is a library for fast approximate clustering and similarity search between dense vectors. It scales to the huge datasets that do not fit in RAM and can be used in a distributed fashion. FAISS is used in our pipeline to perform *k*-means clustering of PQ-quantizated sentence vectors to generate *k*-nearest neighbor edges for similar sentences (nodes) in knowledge network.

**PTBG** [21] (stands for PyTorch BigGraph) is a high-performance graph embedding system allowing distributed training. It was designed to handle large heterogeneous networks containing hundreds of millions of nodes of different types and billions of typed edges. Distributed training is achieved by computing embeddings on disjoint node sets.

### AllenNLP Open Information Extraction

AllenNLP [11] is a powerful library developed by AllenAI that uses PyTorch backend to provide deep-learning models for various natural processing tasks. Specifically, AllenNLP Open Information Extraction provides a trained deep bi-LSTM model for extracting predicates from unstructured text. An API is provided for running inference in both single sentence and batch modes.

## 3 PIPELINE SUMMARY

We briefly summarize the AGATHA semantic graph construction pipeline. It is described in greater detail in the original paper [37].

### Text pre-processing

The input for our system is *a corpora of scientific citations* from the MEDLINE and CORD-19 datasets. These files contain titles and abstracts for millions of biomedical papers. We filter non-English documents, using the FastText Langauge Identification model [16] if the language is not provided. After that we split all abstracts into sentences and process all sentences with ScispaCy library. From each sentence we extract POS-annotated lemmas, entities and perform *n*-gram mining, where *n* ∈ [2, 3, 4] and *n*-grams are composed of frequently co-occurring lemmas. Additionally, we associate all sentences with any relevant metadata, such as the MeSH/UMLS keywords provided along with the citation.

### Semantic Graph Construction

We construct a semantic graph containing different types of nodes, namely, sentences, entities, coded terms (from UMLS and MeSH), *n*-grams, lemmas, and predicates following the schema depicted in Figure 2. Edges between sentences are induced from the nearest-neighbors network of sentence embeddings. We also include an edge between two sentences that appear sequentially within the same abstract, counting the title as the first sentence. Other edges can be inferred directly from the recorded metadata. For instance, the node representing the entity “COVID-19” is connected to every sentence and predicate that discuss COVID-19.

**Figure 1:**
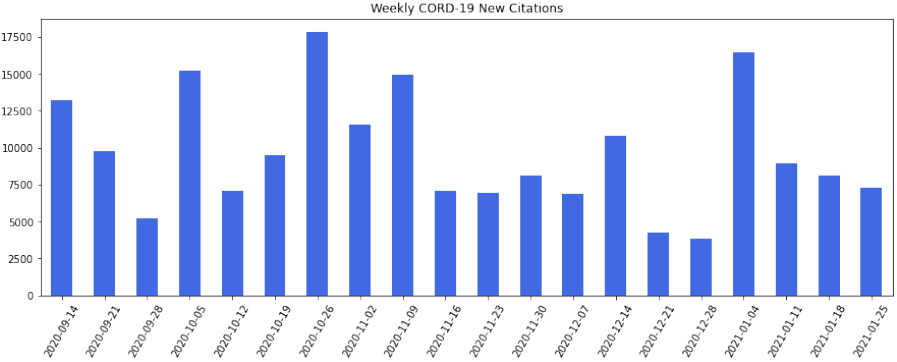
Number of new citations per week in CORD-19 dataset.

**Figure 2:**
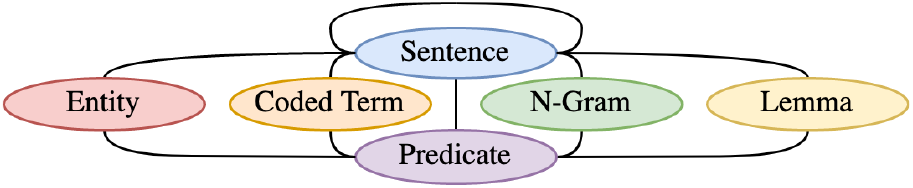
AGATHA multi-layered graph schema.

### NLM UMLS implementation

The prior AGATHA semantic network only includes UMLS terms that appear in SemMedDB predicates [18] which is a major limitation. In this work we enrich the “Coded Term” layer by introducing an additional preprocessing phase wherein we run the SemRep tool with full-fielded output option ourselves *on the entire input corpora*. This phase would be necessary as CORD-19 and most recent MEDLINE citations are not represented within slowly updated SemMedDB. However, we find that we can substantially increase the quality of recovered terms by applying these tools ourselves.

By doing that we not only enrich the “Coded Terms” semantic network layer, but also introduce a significant number of uncovered previously semantic predicates. It happens because SemMedDB is a cumulative database, having various citations in the database processed over many years with various versions of SemRep and various UMLS releases available at different time periods.

To illustrate what was just said, let us consider the following example (PMID: 20109154): *“The results showed that V. cholerae O395 and also other related enteric pathogens have the essential CASS components (CRISPR and cas genes) to mediate a RNAi-like pathway*.*”* The current SemRep version extracts the following predicate: CRISPR-AFFECTS-RNAi, while SemMedDB does not contain any predicates for this sentence. The year of publication of the corresponding paper is 2009, but CRISPR term (C3658200) did not exist in the UMLS metathesaurus on or before 2012, that is why at the time of adding this citation to SemmedDB CRISPR-involved relation could not be identified.

### Graph Embedding

We embed our large semantic graph using a heterogeneous technique that captures node similarity through a *biased transformed dot product*. By explicitly including a bias term for each node, we capture a concepts overall affinity within the network that is critical for such general terms as “coronavirus.” By learning transformations between each pair of node types (e.g., between sentences and lemmas), we enable each type to occupy embedding spaces with differing characteristics. Specifically, we fit an embedding model that optimizes the following similarity measure:

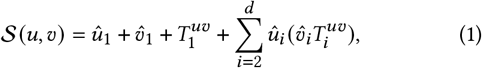

where *u, v* are nodes in the semantic graph with embeddingsû,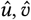, and *T*^*uv*^ is the directional transformation vector between nodes of *u*’s type to nodes of *v*’s.

We use the PTBG heterogeneous graph embedding library to learn *d* = 512 dimensional embeddings for each node of our large semantic graph. While fitting embeddings (*û*) and transformation vectors (*T*^*uv*^), we represent each edge of the semantic graph as two directed edges. These learned values are optimized using softmax loss, where the similarity for one edge is compared against the similarities of 100 negative samples.

### Ranking Semantic Predicates (Transformer model)

After we obtain embeddings per node in the semantic graph, we train AGATHA system ranking model. This model is trained to rank published subject-object pairs above randomly composed pairs of UMLS concepts (negative samples). Two coded terms, along with a fixed-size random subsample of predicates containing each term are input to this model. Graph embeddings for each term and predicate are fed into stacked transformer encoder layers, which apply multi-headed self-attention across the embedding set. The last set of encodings are averaged and the result is projected to the unit interval, forming a scalar prediction for the input’s “plausibility.”

Formally, the model to evaluate term pairs is defined as:

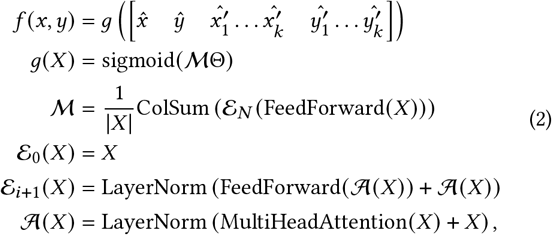

where each *x* ′ and *y*′ are randomly sampled from the neighborhoods of *x* and *y* respectively, and each ^ denotes the graph embedding of the given node. Furthermore, Θ represents a free parameter, which is fit along with parameters internal to each FeedForward and MultiHeadAttention layer, following the standard conventions for each.

The above model is fit using margin ranking loss, where predicates from the training set are compared against a large set of negative samples. Additional details pertaining to specific optimization choices surrounding this model are present in the work originally proposing this model [37].

## 4 AUGMENTING SEMANTIC PREDICATES WITH DEEP LEARNING

We used `SemRep` predicate extraction system in the first system, AGATHA-C, to extract predicates from the abstracts. However, `SemRep` relies on expert coded rules and heuristics to extract biomedical relations leading to significantly fewer predicates for training. Thus, in order to augment the predicates (for the second system, AGATHA-GP) we decided to use a deep learning based information extraction system by Stanvosky *et al*. [31]. Figure 3 shows our overall predicate extraction pipeline.

**Figure 3:**
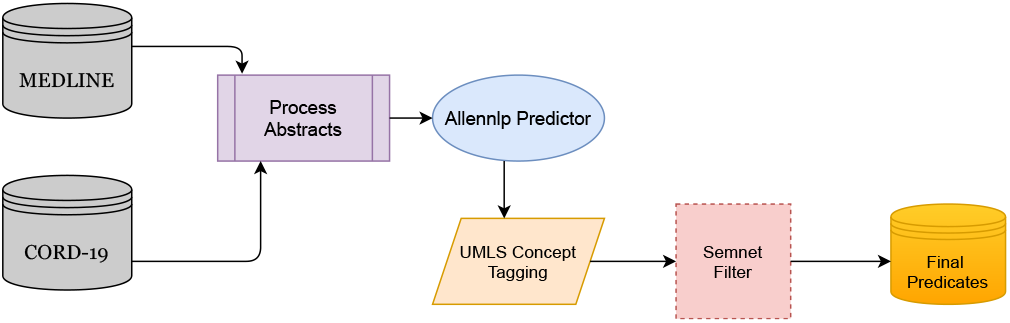
Predicate Extraction pipeline with Deep Learning based Open IE system.

### Abstract Pre-processing

The input for the proposed semantic predicate extraction system is the output files generated by `SemRep` tool with full-fielded output option enabled, obtained from the preprocessing stage described in Sec. 3. As it was mentioned previously, `SemRep` system extracts not only semantic triples, but also maps entities found in the input corpus to their corresponding UMLS concept IDs, this is the data which is used for the following method. The initial set of records includes the sentence raw texts and extracted from them UMLS terms and is augmented throughout the pipeline making it easier to extract final predicates for downstream training.

### Raw Predicate Extraction

We use a pre-trained instance of `RnnOIE` [31] provided as an API by AllenNLP. The model was trained on the `OIE2016` corpus. At a high level the model aims to learn a joint embedding of individual words and their corresponding Beginning-Input-Output (BIO) tags. The output of the model is a probability distribution over the BIO tags. During inference the model selects specific phrases and groups them into `ARG0, V, ARG1` tags. By convention, we treat `ARG0` as the subject and ARG1 as the object in a subject-verb-object tuple. To speed up processing and scale it to thousands of abstracts, we leverage model-parallelism across different machines and run batch-mode inference on chunks of abstracts. Once the model predictions have been extracted we extract the phrases with relevant tags into raw predicates and augment them in the record. A subsequent filtering is performed by extracting the terms matching with previously detected UMLS concepts in the sentence.

### Semnet Filtering

Using a general purpose `RnnOIE` model has it’s own challenges. During processing we noted that a lot of raw predicates were either too general or contained too little meaning to be useful for training a prediction model. To overcome this challenge we designed a corrective filter to reduce noise and retain most useful predicates. We call this filter the *semnet filter*.

Each UMLS concept has an associated semantic type (e.g., COVID-19 has an associated semantic type of `dsyn` (disease)). This is useful for summarizing large set of diverse text concepts into smaller number of categories. We used the metadata from semantic types to construct two networks -a semantic network and a hierarchical network. The semantic network consists of semantic types as nodes and the edges imply a corresponding direct relation between them. The hierarchical network is a network of a semantic type connected to its more general semantic types. For example, a semantic type `dsyn` (disease) is more generally associated with a `biof` (biological function) or a`pathf` (pathological function). In order to filter a predicate, all edges emanating from the subject’s semantic types are computed on a per-predicate basis. These edges also include any specific-general concept relationships. If the object’s semantic type is found to be in the candidate edge set, then we deem the predicate as valid. In our experiments, we found that this filtering method significantly eliminates predicates which do not directly pertain to the biomedical domain.

### Processing Abstracts at Scale

Building a pipeline that scales to thousands of abstracts is not a trivial task. In order to extract predicates from RnnOIE model and extract quality terms of interest we not only have to contend with the problem of running inference on a deep neural network but also the task of *aligning* the extracted terms with the entities recognized by `SemRep`.

#### Deployment details

The RnnOIE model by Stanovsky *et al*. uses a deep Bi-LSTM [27] model to learn the joint word embedding and predict the resulting semantic position tags. Since LSTMs are inherently sequential model, it means that the inference time per sentence would be considerable. We first tried processing an entire collection of abstracts at once on a cluster of 10 machines each consisting of 24 CPUs using the `Dask` [26] library. The entire process took more than 8 hours. Considering that we had about 100 such collections, this inference time was prohibitively high. In order to speed up inference we read each collection once and distributed chunks of abstracts over the machines. This change helped us to cut down the processing time from over a week to just over 4 days for the MEDLINE corpus. For the CORD-19 corpus the processing time was even faster at 2 days. The next step was to align the extracted predicates with the `SemRep` recognized biomedical concepts. We achieved this alignment by first building an index of files that contained a specific abstract ID and then processing the RnnOIE predicates with the aforementioned index. We further optimized the indexing phase by updating the existing index each time we processed more than *τ* abstracts.

The semnet filter does not introduce additional computational overhead and can process a thousand abstracts in under 1 second. Hence, to obtain the most relevant set of predicates we were able to parallelize over “checkpoints” (each of which contained 30k abstracts) in an hour.

## 5 VALIDATION

A fair validation of HG systems is extremely challenging, as these models are designed to predict *novel* connections that are unknown to even those who evaluate the system [34]. In addition, even if validated by rediscovering findings using historical, the process is computationally expensive because of the need to train multiple models to understand how many months (or years) back, the HG system can predict the findings which requires careful filtering of the used papers, vocabulary and other types of data. To present our results in terms of its usefulness for urgent CORD-19-related HG, we use a historical benchmark, which is conceptually described in [37]. This technique is fully automated and does not require any domain experts intervention.

### Positive samples collection

We use `SemRep` and proposed in Sec. 4 approach to process the most recent CORD-19 citations, which were published after the specific cut date making sure that the citations are not included in the training set. After that we extract all subject-object pairs from the obtained results and explicitly check that none of these pairs are presented in the training set. Pairs mentioned in the CORD-19 less than twice are filtered out from the validation set. Almost all of them are either noisy or represent information that already appears in other pairs (e.g., because of the difference in grammar).

We also use the strategy of **subdomain recommendation**. This strategy works in the following way. For each UMLS term we collect its semantic type (which is a part of the metadata provided in UMLS metathesaurus) and group all extracted SemRep pairs by the term-pair criteria (combination of subject and object types). Then we identify the top-20 most common term-pairs subdomains and construct the validation set from pairs belonging to these 20 subdomains.

### Negative samples generation

To generate negative samples per domain, the random sampling is used, that is, for each positive sample we keep its subject and randomly sample the object belonging to the same semantic type as the object of the source pair. We do this 10 times, thus having 10 negative domain-specific samples for each positive sample. When the validation set is generated, we apply our ranking criteria to it, obtaining a numerical score value *s* per each sample, where *s* ∈ [0, 1].

### Evaluation metrics

We propose our approach as a recommendation system and to report our results we use a combination of the following classification and recommendation metrics.

- Classification metrics: (1) Area under the receiver-operating-characteristic curve (AUC ROC); (2) Area under the precision-recall curve (AUC PR).
- Recommendation metrics: (1) Top-k precision (P.@k); (2) Average precision (AP.@k); and (3) Overall reciprocal rank (RR).

We report these numbers in per subdomain manner to better understand how the system performs with respect to specific task (e.g. drug repurposing).

## 6 RESULTS

To report results, we provide the performance measures for three AGATHA models trained on the same input data (MEDLINE corpus and CORD-19 abstracts dataset):

1. AGATHA-O: Baseline AGATHA model [37];
2. AGATHA-C: AGATHA-O with new UMLS layer and SemRep enrichment;
3. AGATHA-GP: AGATHA-C with additional deep learning-based extracted and further filtered predicates.

It is done in this particular manner because the major role in learning the proposed ranking criteria depends heavily on the quality of extracted semantic predicates and their number, as they form the training set for the AGATHA ranking module. At the moment of writing, no other general purpose and available for public use HG system compliant with the three validation criteria, namely, (a) ability to run thousands of queries in a reasonable time, (b) ability to process COVID-19 related vocabulary, and (c) ability to operate in multiple domains was available for comparison.

The performance of both AGATHA-C and AGATHA-GP allows to run thousands of queries in a very short time (in the order of minutes), making the validation on a large number of samples possible. Unfortunately, given the current circumstances, large-scale validation for the specific scientific subdomain (COVID-19 related hypotheses) is hard to implement, because well-established and reliable factual base is being actively developed at the moment and big historic gap for the vocabulary simply does not exist (e.g., the COVID-19 term is just approximately one year old). We, however, provide the validation set including 2736 positive connections extracted from CORD-19 dataset citations added within the time frame from October 28, 2020 to January 21, 2021, which numbered at 77 thousand abstracts.

In Table 1, we share some basic graph metrics for the models AGATHA-O, AGATHA-C and AGATHA-GP. The most significant change is observed in the number of semantic predicates and coded terms, which clearly represents the purpose of introducing additional preprocessing steps.

**Table 1:** Graph metrics (M = millions, B = billions).

| Node Type | Counts |  |  |
| --- | --- | --- | --- |
|  | AGATHA-O | AGATHA-C | AGATHA-GP |
| Sentence | 190.6 M. | 190.6 M. | 190.6 M. |
| Predicate | 24.2 M. | 36.3 M. | 38.7 M. |
| Lemma | 16.8 M. | 16.1 M. | 16.1 M. |
| Entity | 41.7 M. | 43.2 M. | 43.2 M. |
| Coded Term | 538,588 | 855,351 | 855,351 |
| *n*-Grams | 212.922 | 326.864 | 333.575 |
| Total Nodes | 274.1 M. | 287.4 M. | 289.8 M. |
| Total Edges | 13.52 B. | 13.5 B. | 13.53 B. |

In Table 2, we compare aforementioned models using the metrics described in Sec. 5. We present predicate types with NLM semantic type codes [23] due to space restrictions. *Both AGATHA-C and AGATHA-GP models show significant gains when compared to AGATHA-O baseline model*. Benefits in the most problematic for the baseline model areas (e.g., *(Gene)* → *(Gene)* denoted by *(gngm,gngm)*) serve the best illustration for that, showing up to almost 30 percent advantage in ROC AUC. Now all most popular biomedical subdomains are covered by the proposed models and show AUC ROC results at at least 0.87. Average ROC AUC value is increased by 0.09.

**Table 2:**
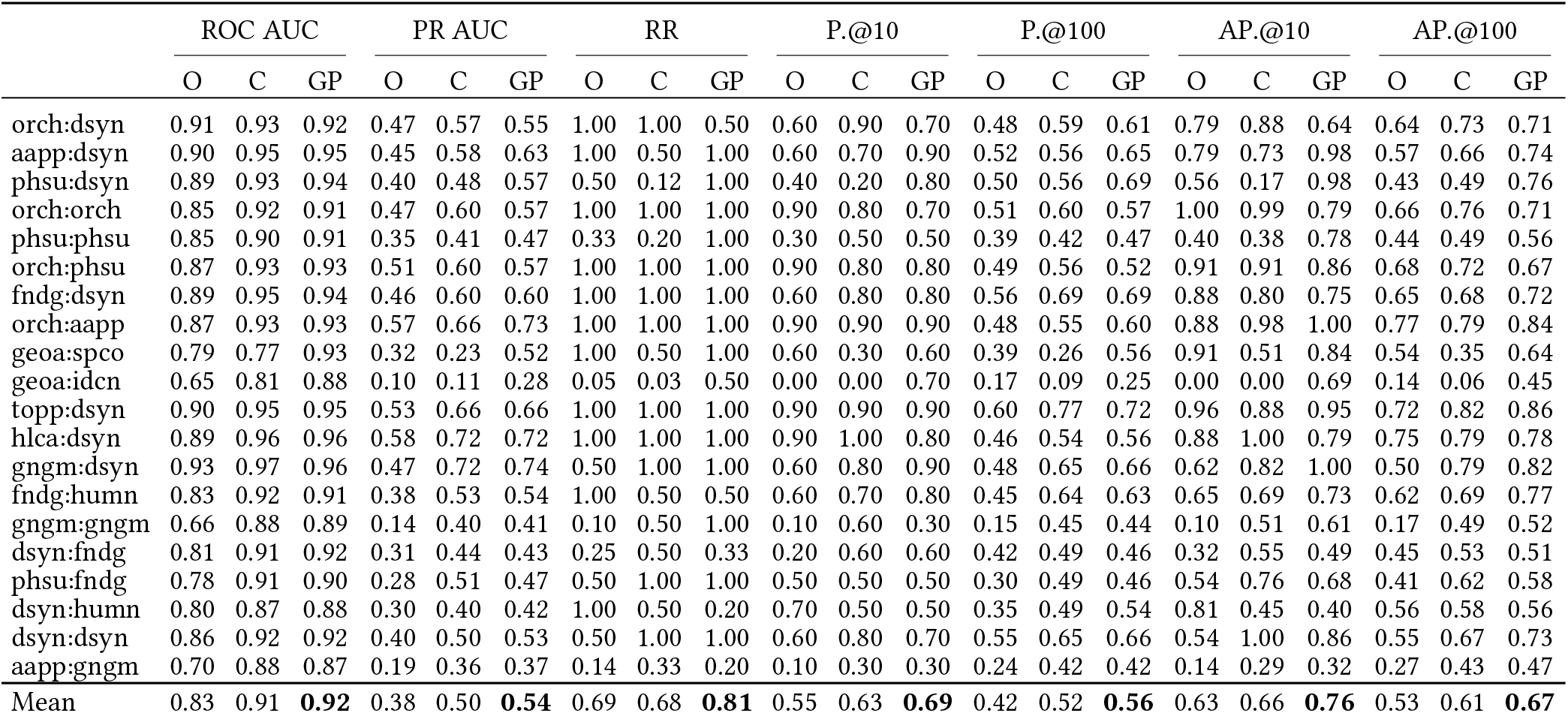
Classification and recommendation quality metrics across recently popular COVID-19-related biomedical subdomains. Labels O, C and GP stand for AGATHA-O, AGATHA-C and AGATHA-GP models, respectively.

|  | ROC AUC |  |  | PR AUC |  |  | RR |  |  | P.@10 |  |  | P.@100 |  |  | AP.@10 |  |  | AP.@100 |  |  |
| --- | --- | --- | --- | --- | --- | --- | --- | --- | --- | --- | --- | --- | --- | --- | --- | --- | --- | --- | --- | --- | --- |
|  | O | C | GP | O | C | GP | O | C | GP | O | C | GP | O | C | GP | O | C | GP | O | C | GP |
| orch:dsyn | 0.91 | 0.93 | 0.92 | 0.47 | 0.57 | 0.55 | 1.00 | 1.00 | 0.50 | 0.60 | 0.90 | 0.70 | 0.48 | 0.59 | 0.61 | 0.79 | 0.88 | 0.64 | 0.64 | 0.73 | 0.71 |
| aapp:dsyn | 0.90 | 0.95 | 0.95 | 0.45 | 0.58 | 0.63 | 1.00 | 0.50 | 1.00 | 0.60 | 0.70 | 0.90 | 0.52 | 0.56 | 0.65 | 0.79 | 0.73 | 0.98 | 0.57 | 0.66 | 0.74 |
| phsu:dsyn | 0.89 | 0.93 | 0.94 | 0.40 | 0.48 | 0.57 | 0.50 | 0.12 | 1.00 | 0.40 | 0.20 | 0.80 | 0.50 | 0.56 | 0.69 | 0.56 | 0.17 | 0.98 | 0.43 | 0.49 | 0.76 |
| orch:orch | 0.85 | 0.92 | 0.91 | 0.47 | 0.60 | 0.57 | 1.00 | 1.00 | 1.00 | 0.90 | 0.80 | 0.70 | 0.51 | 0.60 | 0.57 | 1.00 | 0.99 | 0.79 | 0.66 | 0.76 | 0.71 |
| phsu:phsu | 0.85 | 0.90 | 0.91 | 0.35 | 0.41 | 0.47 | 0.33 | 0.20 | 1.00 | 0.30 | 0.50 | 0.50 | 0.39 | 0.42 | 0.47 | 0.40 | 0.38 | 0.78 | 0.44 | 0.49 | 0.56 |
| orch:phsu | 0.87 | 0.93 | 0.93 | 0.51 | 0.60 | 0.57 | 1.00 | 1.00 | 1.00 | 0.90 | 0.80 | 0.80 | 0.49 | 0.56 | 0.52 | 0.91 | 0.91 | 0.86 | 0.68 | 0.72 | 0.67 |
| fndg:dsyn | 0.89 | 0.95 | 0.94 | 0.46 | 0.60 | 0.60 | 1.00 | 1.00 | 1.00 | 0.60 | 0.80 | 0.80 | 0.56 | 0.69 | 0.69 | 0.88 | 0.80 | 0.75 | 0.65 | 0.68 | 0.72 |
| orch:aapp | 0.87 | 0.93 | 0.93 | 0.57 | 0.66 | 0.73 | 1.00 | 1.00 | 1.00 | 0.90 | 0.90 | 0.90 | 0.48 | 0.55 | 0.60 | 0.88 | 0.98 | 1.00 | 0.77 | 0.79 | 0.84 |
| geoa:spco | 0.79 | 0.77 | 0.93 | 0.32 | 0.23 | 0.52 | 1.00 | 0.50 | 1.00 | 0.60 | 0.30 | 0.60 | 0.39 | 0.26 | 0.56 | 0.91 | 0.51 | 0.84 | 0.54 | 0.35 | 0.64 |
| geoa:icdn | 0.65 | 0.81 | 0.88 | 0.10 | 0.11 | 0.28 | 0.05 | 0.03 | 0.50 | 0.00 | 0.00 | 0.70 | 0.17 | 0.09 | 0.25 | 0.00 | 0.00 | 0.69 | 0.14 | 0.06 | 0.45 |
| topp:dsyn | 0.90 | 0.95 | 0.95 | 0.53 | 0.66 | 0.66 | 1.00 | 1.00 | 1.00 | 0.90 | 0.90 | 0.90 | 0.60 | 0.77 | 0.72 | 0.96 | 0.88 | 0.95 | 0.72 | 0.82 | 0.86 |
| hlca:dsyn | 0.89 | 0.96 | 0.96 | 0.58 | 0.72 | 0.72 | 1.00 | 1.00 | 1.00 | 0.90 | 1.00 | 0.80 | 0.46 | 0.54 | 0.56 | 0.88 | 1.00 | 0.79 | 0.75 | 0.79 | 0.78 |
| gngm:dsyn | 0.93 | 0.97 | 0.96 | 0.47 | 0.72 | 0.74 | 0.50 | 1.00 | 1.00 | 0.60 | 0.80 | 0.90 | 0.48 | 0.65 | 0.66 | 0.62 | 0.82 | 1.00 | 0.50 | 0.79 | 0.82 |
| fndg:humn | 0.83 | 0.92 | 0.91 | 0.38 | 0.53 | 0.54 | 1.00 | 0.50 | 0.50 | 0.60 | 0.70 | 0.80 | 0.45 | 0.64 | 0.63 | 0.65 | 0.69 | 0.73 | 0.62 | 0.69 | 0.77 |
| gngm:gngm | 0.66 | 0.88 | 0.89 | 0.14 | 0.40 | 0.41 | 0.10 | 0.50 | 1.00 | 0.10 | 0.60 | 0.30 | 0.15 | 0.45 | 0.44 | 0.10 | 0.51 | 0.61 | 0.17 | 0.49 | 0.52 |
| dsyn:fndg | 0.81 | 0.91 | 0.92 | 0.31 | 0.44 | 0.43 | 0.25 | 0.50 | 0.33 | 0.20 | 0.60 | 0.60 | 0.42 | 0.49 | 0.46 | 0.32 | 0.55 | 0.49 | 0.45 | 0.53 | 0.51 |
| phsu:fndg | 0.78 | 0.91 | 0.90 | 0.28 | 0.51 | 0.47 | 0.50 | 1.00 | 1.00 | 0.50 | 0.50 | 0.50 | 0.30 | 0.49 | 0.46 | 0.54 | 0.76 | 0.68 | 0.41 | 0.62 | 0.58 |
| dsyn:humn | 0.80 | 0.87 | 0.88 | 0.30 | 0.40 | 0.42 | 1.00 | 0.50 | 0.20 | 0.70 | 0.50 | 0.50 | 0.35 | 0.49 | 0.54 | 0.81 | 0.45 | 0.40 | 0.56 | 0.58 | 0.56 |
| dsyn:dsyn | 0.86 | 0.92 | 0.92 | 0.40 | 0.50 | 0.53 | 0.50 | 1.00 | 1.00 | 0.60 | 0.80 | 0.70 | 0.55 | 0.65 | 0.66 | 0.54 | 1.00 | 0.86 | 0.55 | 0.67 | 0.73 |
| aapp:gngm | 0.70 | 0.88 | 0.87 | 0.19 | 0.36 | 0.37 | 0.14 | 0.33 | 0.20 | 0.10 | 0.30 | 0.30 | 0.24 | 0.42 | 0.42 | 0.14 | 0.29 | 0.32 | 0.27 | 0.43 | 0.47 |
| Mean | 0.83 | 0.91 | **0.92** | 0.38 | 0.50 | **0.54** | 0.69 | 0.68 | **0.81** | 0.55 | 0.63 | **0.69** | 0.42 | 0.52 | **0.56** | 0.63 | 0.66 | **0.76** | 0.53 | 0.61 | **0.67** |

Our validation strategy involves a big number of many-to-many queries, making the area under precision-recall curve another very illustrative metric. This is where the newly proposed models show even more drastic improvements over the baseline AGATHA-O. For some subdomains, like *(Gene or Genome)* → *(Gene or Genome) (gngm,gngm)* or *(Amino Acid, Peptide, or Protein)* → *(Gene or Genome) (aapp,gngm)*, we observe that new models take the recommendations performance to the new quality level. Average PR AUC value is increased by 0.16.

The approximate running time with corresponding types of used hardware is presented in Table 3. Each row corresponds to the stage in the AGATHA-C /AGATHA-GP pipelines. The column “M” (machines) and CPU show the number of machines and required CPUs, respectively. In the column “GPU” we indicate if GPU was required or optional. For AGATHA training we used two NVIDIA V100 per machine. The minimal requirements for RAM per machine are in column “RAM”. The running time of queries is negligible.

**Table 3:** Running time and hardware requirements.

| Stage | Time | Hardware |  |  |  |
| --- | --- | --- | --- | --- | --- |
|  |  | M | CPU | GPU | RAM |
| SemRep Processing | 2 d | 10-28 | 20+ | Opt | N/A |
| AllenNLP Predicates | 3 d | 28-40 | 20+ | Opt | N/A |
| Graph Construction | 10 d | 30+ | 20+ | Opt | 120GB+ |
| Graph Conversion | 7 h | 1 | 40+ | Opt | 1TB+ |
| Graph Embedding | 1 d | 20 | 24+ | Opt | 120GB+ |
| AGATHA Training | 22 h | 5+ | 2+ | Yes | 300GB+ |
| Network Adjacency | 1 d | 1 | 40+ | Opt | 1.5TB+ |

## 7 CASE STUDY

The proactive discovery of ongoing research findings is an important component in the validation of hypothesis generation systems [36]. In particular, in the current uncertain situation when a lot of unintentionally incorrect discoveries are published, the validation must include human-in-the-loop part even in limited capacity such as in [2, 30]. To demonstrate the predictive potential of AGATHA-C we perform a case study on three COVID-19-related novel connections manually selected by the domain expert. These connections were published after the cut date before which any data used in training was available to download at NIH.

At a low level, all AGATHA models use entity subsampling to calculate pairwise ranking criteria, which means that the absolute numbers may fluctuate slightly. Thus, to present the numeric scores, each experiment was repeated 100 times to compute the average and standard deviation that we present in Table 4.

**Table 4:** Scores for valid recently published connections obtained by different AGATHA models. Reported average values for 100 runs and standard deviation.

|  | AGATHA-O | AGATHA-C | AGATHA-GP |
| --- | --- | --- | --- |
| COVID-19:Melatonin | 0.63 ± 0.03 | 0.91 ± 0.03 | 0.78 ± 0.03 |
| COVID-19:Oxytocin | 0.75 ± 0.03 | 0.98 ± 0.02 | 0.81 ± 0.02 |
| COVID-19:BST2 gene | 0.41 ± 0.01 | 0.88 ± 0.03 | 0.74 ± 0.03 |

AGATHA-C was tested whether it will be able to predict compounds potentially applicable for the treatment of COVID-19 and the genes involved in the SARS-CoV-2 pathogenesis. The data confirming cardiovascular protective effects of hormone oxytocine were published recently [9, 40]. The protective effect is linked to anti inflammatory activity of the hormone. For this connection AGATHA-C generated the score of 0.98.

Similarly, we tested the prediction of the effects of the other hormone, melatonin. Several publications, started from November 2020 [3, 8, 13, 43] show the protective effects of melatonin, specifically for COVID-19 neurological complications. The activity was linked to anti-oxidative effects of the melatonin. For this connection AGATHA-C generated the score of 0.91.

Our system accurately predicted with score of 0.88 the involvement of tetherin (BST2). The results published in 2021 [32] show that tetherin restricts the secretion of SARS-CoV-2 viral particles and is downregulated by SARS-CoV-2. Therefore, pharmacological activation of tetherin expression, or inhibition of the degradation could be a promising direction of the development of SARS-CoV-2 treatment.

## 8 LESSONS LEARNED AND OPEN PROBLEMS

### Quality of the information retrieval pipelines

Information retrieval is an important part of any HG pipeline. In order to uncover *implicit* connections, the system should be able to capture existing *explicit* connections with as much quality as possible. Given that human knowledge is usually stored in a non-structured manner (e.g., scientific texts), the quality of systems that process raw textual data, such as those that solve the named entity recognition, or word sense disambiguation problems, is crucial.

We observed that the SemRep system performs better concept and relation recognition when full abstracts are used as input data instead of single sentences. SemRep also allows to perform optional sortal anaphora resolution to extract co-references to the entities from neighbouring sentences, which was shown to be useful in [17] and is used in this work.

### “Positive” research bias

The absence of published negative research results is a big problem for the HG field. With mostly positive results available, often we have to generate negative examples through some kind of random sampling. These negative samples likely do not adequately represent the real nature of negatively confirmed scientific findings. Likely, one of the most important future work directions in the area of HG is to accurately distinguish and leverage positive and negative proposed results.

### Domain experts involvement

When any hypothesis generation system is built, one of the first questions a designer should address is extent that domain experts are expected to participate in the pipeline. Modern decision-making systems allow a fully automated discovery process (like the AGATHA system), but this may not be sufficient. A domain expert who interfaces with a HG system as a black box may not trust generated results or know how best to interpret them. The challenge of interpretable hypothesis generation remains a significant barrier to widespread adoption of these kinds of research tools. For this we advocate using our “structural” learning HG system MOLIERE [35] in which with the topical modeling and network analytic measures we interpret and explain the results.

### The nature of input corpora

The question of what should be used as input to a topic-modeling based hypothesis generation system is raised in [34]. Using full-text papers shows an improvement, but the trade-off between run time and output quality was barely justifiable. However, deep learning models have a greater potential for extracting useful information from large input sources, and as it was demonstrated in our previous work [37], show significant performance advancements. Thus the question of using full-text papers in deep learning-based hypothesis generation systems should be addressed. Unfortunately, it is currently too computationally expensive our resources as the number of sentences and thus predicates and edges will be significantly larger.

### Knowledge resolution

Our newly proposed systems showed that the knowledge resolution plays a major role in subdomain recommendation. To increase the scope of model expertise (and the scope of potential applications beyond the biomedical fields) we deliberately incorporate a general-purpose information retrieval system RnnOIE into AGATHA-GP. This additional information results in significant gains in broad subdomains like *(Geographic Area)* → *(Idea or Concept) (geoa,idcn)*. At the same time, we observe that AGATHA-C performs better in “microscopic” biomedical areas, e.g. *(Organic Chemical)* → *(Organic Chemical) (orch,orch)*, which raises the question of choosing the appropriate model for every specific use case. Although, both systems process all types of queries, the general purpose predicates participated in training significantly improve “macroscopic” types of queries.

## 9 RELATED WORK

A number of works have been proposed to organize the CORD-19 literature into a structured knowledge graph for different purposes. For instance, Basu *et al*. [5] propose ERLKG -a knowledge graph built on CORD-19 with entities corresponding to gene/chemical/disease names and the edges forming relations between the concept. They use a fine tuned SciBERT model for both entity and relation extraction. The main purpose of the knowledge graph is to predict a link between a given chemical-disease and chemical-protein pair using a trained GCN autoencoder [19] approach. In another similar work, Oniani *et al*. [25] build a co-occurrence network on a subset of CORD-19 with the edges corresponding to either gene-disease, gene-mutation or chemical-disease type. The network is then embedded into latent space using a node2vec walk. Link prediction is performed on the nodes by training different classical machine learning algorithms. A major shortcoming of these approaches is that they limit themselves to either specific kind of entities or relations or both and as a result not only the scope of possible new literature is narrowed but a lot of additional useful knowledge is filtered out of the system. In contrast, our system does not limit itself to specific entity or relation type and is able to capture much more information from the same corpus.

A major interest of constructing knowledge graphs is to allow medical researchers to re-purpose existing drugs for treating COVID-19. Zhang *et al*. [42] develop a system that uses combined semantic predications from SemMedDB and CORD-19 (extracted using SemRep) to recommend drugs for COVID-19 treatment. To improve the predications from CORD-19, the authors fine tune various transformer based models on a manually annotated internal dataset. Their resulting knowledge graph consists of 131,555 nodes and 2,558,935 edges. Our work on the other hand utilizes similar technologies and produces a bigger graph with 287,356,836 nodes and 13,500,291,256 edges. Moreover, we do not post-process extracted relations from SemRep and are still able to achieve a higher RoC metric. Another system proposed by Martinc *et al*. [22] uses a fine-tuned SciBERT model to generate contextualized embeddings of CORD-19 articles and using an initial seed set of targets proposes possible therapy targets. However, this system is very different from ours as it treats the entire article as a bag of words and directly trains a word embedding model on CORD-19. It was earlier noted that KinderMiner [20] provides a web-based literature discovery tool and supports COVID-19 queries. The underlying algorithm is based on a simple keyword co-count between source and target words in a given corpus. While co-count is a fast and scalable approach, it suffers from a lack of “discrimination” i.e. two keywords occurring together more frequently do not always imply a high degree of correlation.

The vastness of COVID-19 literature also spurned the need for having systems that could allow researchers and base users alike to get their COVID-19 queries answered. Systems like CKG (Wise *et al*.) [41] and SciSight (Hope *et al*.) [14] currently provide this functionality. While we do aim to provide an easy to use web-framework for medical researchers, the scope of the aforementioned systems is beyond the scope of our work. Unfortunately, no existing system out of those that are trained to accept terms related to COVID-19 or SARS-CoV-2 provided an open access for massive validation for a fair comparison with or was able to be tested in multiple domains like AGATHA-C.

## 10 CONCLUSIONS

We present two graph mining transformer based models AGATHA-C and AGATHA-GP, for micro-and macroscopic scales of queries respectively, which are designed to help domain experts solve high-priority research problems and accelerate scientific discovery. We perform per-subdomain validation of these new models on a rapidly changing COVID-19 focused dataset, composed of recently published concept pairs and demonstrate that the proposed models achieve state-of-the-art prediction quality. Both models significantly outperform the existing baseline system AGATHA-O. We deploy the proposed models to the broad scientific community and believe that our contribution can raise more interest in prospective hypothesis generation applications.

